# BML-257, a small molecule that protects against drug induced liver injury in zebrafish

**DOI:** 10.1101/2022.01.09.475146

**Authors:** Urmila Jagtap, Sandeep Basu, Lavanya Lokhande, Nikhil Bharti, Chetana Sachidanandan

**Author notes:** These authors contributed equally to this work (Lavanya Lokhande, Nikhil Bharti).

## Abstract

The use of many essential drugs is restricted due to their deleterious effects on the liver. Molecules that can prevent or protect the liver from drug induced liver injury (DILI) would be valuable in such situations. We used hepatocyte-specific expression of bacterial nitroreductase in zebrafish to cause temporally controlled liver damage. This transgenic line was used to run a whole organism based chemical screen in zebrafish larvae. In this screen we identified BML-257, a potent small molecule AKT inhibitor, that protected the liver against metronidazole-induced liver injury. BML-257 also showed potent prophylactic and pro-regenerative activity in this liver damage model. BML-257 also showed remarkable protective action in two independent toxicological models of liver injury caused by acetaminophen and Isoniazid. This suggests that BML-257 may have the potential to protect against multiple kinds of drug induced liver injury.

## Introduction

Many drugs cause mild to severe hepatotoxicity and drug induced liver injury (DILI) is a major cause for withdrawal of approved drugs. However, a number of life-saving drugs with hepatotoxic effects continue to be used because of lack of safer alternatives [1-3]. Currently there are no well-established compounds with proven hepatoprotective activity that are clinically accepted. Screens for hepatoprotective compounds have been conducted on cultured hepatocytes identifying candidate molecules [3-6]. However cultured cells cannot model the whole organism in terms of drug processing or exploring the role of non-hepatocytes in liver damage and regeneration. Although rodents have been used extensively to study liver damage and regeneration, these systems are not ideal for screening small molecules. Zebrafish with its complex physiology and well worked out genetics has been very useful in uncovering the pathophysiology of various diseases [7, 8]. Zebrafish has also been at the forefront of high content chemical screening due to the permeability of the zebrafish embryos to small molecules and their visual accessibility [9-12].

Zebrafish has been used very successfully in studying liver development and diseases that affect the liver such as hereditary liver diseases [13], biliary defects [14, 15], fatty liver diseases [16] and hepatocellular carcinoma [17, 18]. We have previously characterized a cell-ablation model for liver damage and regeneration in the adult zebrafish [19]. Drug induced liver injury has been modeled in zebrafish with various drugs such as acetaminophen [20], ibuprofen [21], rifampicin and dexamethasone [22]. Pfizer lnc. And Jansen pharmaceutical companies also validated zebrafish as a valuable model system during their blinded study of identifying comounds that could cause DILI in humans [23]. A chemical screen for small molecule therapeutic agents against acetaminophen (APAP) induced liver injury in the zebrafish embryos identified Prostaglandin E2 (PGE2), which works in synergy with N-acetyl cysteine to protect from APAP-induced liver injury [20].

In this study, we have utilized a transgenic system to cause hepatocyte-specific cell ablation. When the transgenic zebrafish larvae are treated with the prodrug zebrafish it leads to hepatocyte death [24, 25]. The loss followed by regeneration of hepatocytes can be quantified using a liver-specific mCherry fluorescence. We performed a screen for small molecules that prevent the MTZ-induced liver damage and identified a potent compound, BML-257. BML-257, an AKT inhibitor, also showed significant prophylactic action when the liver was pre-exposed to the compound before damage. In addition, the compound was effective in augmenting the regenerative ability of the liver post-damage. We tested BML-257 in two different toxicological models of liver injury using Acetaminophen and Isoniazid and found a significant recovery of the damaged liver in these models. Thus BML-257 shows potential for use as an antidote for drug-induced-liver-injury.

## Materials and methods

### Zebrafish maintenance

Zebrafish (Danio rerio) were kept in 10-liter tanks and bred, raised, and maintained at 28.5°C under standard conditions as described [26]. All zebrafish experiments complied with the standards of the Animal Ethics Committee (IAEC) of the CSIR-Institute of Genomics and Integrative Biology, India in accordance with the recommendations of the Committee for the Purpose of Control and Supervision of Experiments on Animals (CPCSEA), Govt. of India.

### Breeding and embryo collection

Male and female fish were kept in breeding tanks separated by dividers overnight. Day-night cycle (14 hours-light and 10 hours-dark) was strictly maintained. The next day, dividers were removed just before the day cycle started. Fish were allowed to mate and embryos were collected. Embryos were sorted and approximately 100 embryos per petri plate (90mm) were kept at 28°C in the E3 buffer. For all experiments 0.003% PTU was added at 1-day post fertilization (dpf) to avoid melanin pigment formation.

### Zebrafish lines

The double transgenic zebrafish line used in this study was generated by crossing two separate lines. The hepatocyte-specific promoter of fabp10a drives the expression of the Gal4-VP16 transcription activator in the *Tg(fabp10a:GAL4-VP16,myl7:Cerulean)* [24] line. The *Tg(UAS:nfsb-mcherry)* would express nfsb or bacterial nitroreductase fused to mCherry in all the cells where GAL4-VP16 is expressed. Thus, the double transgenic line will have expression of nfsb and mcherry in the hepatocytes [25]. All the experiments were performed in the transgenic zebrafish line.

### Assay for liver damage by Metronidazole

At 3-days post fertilization (dpf) the transgenic zebrafish larvae were treated with freshly prepared Metronidazole (MTZ) in 0.2% DMSO in E3 buffer (5 mM NaCl, 0.17 mM KCl, 0.33 mM CaCl2, 0.33 mM MgSO4). The treatment was given in a 90mm petri plate wherein 100 larvae were exposed to 25ml of MTZ solution for 24 hours in a dark humid chamber and the plate was kept in the incubator at 28°C. After the treatment, MTZ was either washed off for further chemical treatment on the live larvae or fixed in 4% PFA.

### Chemical screening

Kinase inhibitors library (Enzo, SCREEN-WELL®, Cat no. BML-2832-0100) was used for the liver protection assay. A randomly selected pool of 25 larvae (3 dpf) was placed in each well in a 12 well plate. 2 ml of MTZ solution (2.5mM in 0.2% E3 buffer) was added in each well. In the same solution, the desired concentration of the test drug was added. ‘Only DMSO’ treatment served as a control. The plate was incubated at 28°C in a dark humid chamber and after 24 hours, the larvae were fixed in 4% PFA for imaging or RNAiso Plus (Takara, 9109) for RNA isolation.

### Assay for liver regeneration

MTZ treatment to mCherry positive transgenic larvae was given as mentioned in an earlier section (3dpf to 4dpf). After 24 hours of treatment, MTZ was washed off and the larvae were rinsed with E3 water at least 5 times. A pool of randomly selected 25 larvae was placed in 12 well plates with 2ml of E3 buffer in each well. larvae were exposed to 10µM BML-257 from 4dpf to 6dpf (i.e., 48hours) at 28°C in a dark humid chamber. At 6dpf, the larvae were fixed in 4% paraformaldehyde (PFA) for overnight and imaged for mCherry fluorescence under fluorescence microscope (Zeiss Axioscope.A1).

### Prophylactic treatment

3dpf larvae were treated with the drug of interest i.e., DMSO or BML-257 at a desired concentration for 48 hours. At 5dpf, the compound was removed and the larvae were washed with 2ml E3 water twice. MTZ treatment (2.5mM in 0.2% E3 buffer) was given for 24 hours following that the plates were incubated at 28°C in a dark humid chamber. At 6dpf, the larvae were fixed in 4% PFA for imaging.

### Assay for liver protection

A randomly selected pool of 25 larvae (3dpf) was placed in a 12 well plate. Larvae were treated with 2ml of standardized concentration of the liver damaging agent i.e., Acetaminophen (15mM) or Isoniazid (10mM) for 24hours. In the same solution, the desired concentration of BML-257 i.e., 1.25 µM or 2.5µM was added. ‘Only DMSO’ treatment served as a control. The plate was incubated at 28°C in a dark humid chamber and after 24 hours, the larvae were fixed in 4% PFA and imaged for mCherry fluorescence.

### Quantification of fluorescence using ImageJ

Using ImageJ software, the fluorescent liver images were transformed into RGB stack. The red stack was used for measurement of mCherry fluorescence. A fixed sized rectangle covering the whole liver was used for each set. Using the analyze tool, area, area of integrated density and mean grey value for each image were determined. A similar procedure was followed to measure the background fluorescence. The background fluorescence density was subtracted for each larval liver. The average of mean fluorescence density between the control and test sample was plotted as a scatter plot using Graphpad prism software and Student’s *t* test (unpaired) was used for statistical analysis.

### ROS measurement

The assay was performed in whole larvae. After MTZ treatment, the larvae were placed in 96 well transparent flat bottom plate (3 larvae/well). 200µl of 2’,7’ – dichlorofluorescindiacetate (H2DCFDA) (Invitrogen, ab113851) was added in each well and the plate was incubated at 37°C in the dark. After 30 minutes, the fluorescence was measured at 485/535nm on multimode plate reader Tecan-infinite M200PRO every 15 minutes for 25 cycles. 0.1µl H_2_O_2_ served as a positive control. Background fluorescence was measured and was subtracted from the respective test samples.

### Real time quantitative PCR

A total of 1µg RNA was reverse transcribed using a QuantiTect Reverse Transcription kit (Qiagen, 205313). The amplification efficiency of primers was determined by PCR amplification with serial dilutions of cDNA. Only primer pairs with 100% amplification efficiency were used. Primers used for amplification: fabp10a forward:CACCTCCAAAACTCCTGGAA and reverse:TTCTGCAGACCAGCTTTCCT and rpl13a forward:TCTGGAGGACTGTAAGAGGTATGC and reverse: AGACGCACAATCTTGAGAGCAG. RT-qPCR was subsequently performed as described [23] using Sybr green (Roche, 06924204001) on The *LightCycler* 480 System. All genes were normalized against *rpl13a*. Analyses on normalized data were performed using the 2^−ΔΔ^ CT algorithm. Three independent experiments were conducted for each condition. Student’s t test was used to calculate statistical significance of the data.

### Western Blot Analysis

After respective treatments, the larvae were collected and homogenized in the NP40 lysis buffer (ThermoFisher, FNN0021). The homogenized liver lysates were centrifuged (14,000 rpm for 20 min), and supernatants were separated for the assay. BCA Protein Assay Kit (Pierce 23225) was used for estimation of protein concentration. 40μg of each sample was run on 10% SDS-Polyacrylamide gel. Following transfer of the proteins onto a PVDF membrane (Merck Milipore, ISEQ00010), the blots were saturated with 5% BSA in Tris-buffered saline (TBS) containing 0.1% (v/v) Tween-20 (TBST) for 2h and probed overnight with 1:5000 diluted primary antibody (Abcam, ab167453). Unbound primary antibody was removed by washing the membrane with TBST thrice. Following this, the blots were incubated with 1:10000 diluted HRP conjugated antirabbit IgG (Cell Signaling, 7074S) for 2h. The peroxidase signal was detected using EMD Millipore Immobilon Western Chemiluminescent HRP Substrate (WBKLS0500).

### Whole Mount RNA in Situ Hybridization

Larvae were fixed in 4% paraformaldehyde overnight. Larvae were subjected to methanol fixation. Prior to RNA in situ hybridization the larvae were rehydrated through a series of methanol dilutions. 3 dpf larvae were treated with 20 μg/mL proteinase K for 13 min followed by the fixation with 4% PFA. RNA in situ hybridization was performed as described [27]. A Zeiss (Stemi 2000CVR) bright field microscope was used to capture images at 5X magnification (with AxiocamICc1).

## Results and Discussion

### Metronidazole induced hepatocyte damage and regeneration in zebrafish larvae

We have used a genetic cell ablation system in zebrafish to cause liver injury [28]. The double transgenic line *Tg(fabp10a:GAL4-VP16,myl7:Cerulean)::(UAS:nfsb-mCherry)* [24, 25] expresses the bacterial nitroreductase (*nfsb*)-mCherry fusion protein in a specifically in the hepatocytes (Fig. 1A) (hitherto known as *fabp10a-nfsb*). Treatment of zebrafish larvae with Metronidazole (MTZ) causes hepatocyte-specific cell death. The bacterial *nfsb* reduces the pro-drug MTZ into a toxic product leading to elevated ROS levels [29]. Since zebrafish larval liver is functional and the fabp10a promoter is fully active by 3 days post-fertilization (dpf) [30] we exposed larvae to various concentrations of MTZ (10mM, 5mM, 2.5mM and 1.25mM) for 24 hours. The larval liver was imaged for mCherry fluorescence (Fig. 1B, upper panel) and the fluorescence was quantified using ImageJ software. All concentrations of MTZ led to significant loss of mCherry fluorescence indicating damage (Fig. 1B). However, 10mM and 5mM MTZ also lead to larval death and morbidity. 2.5mM MTZ caused damage without noticeable effect on the health of the larvae. Thus, we chose 2.5mM MTZ for all further experiments.

**Figure 1:**
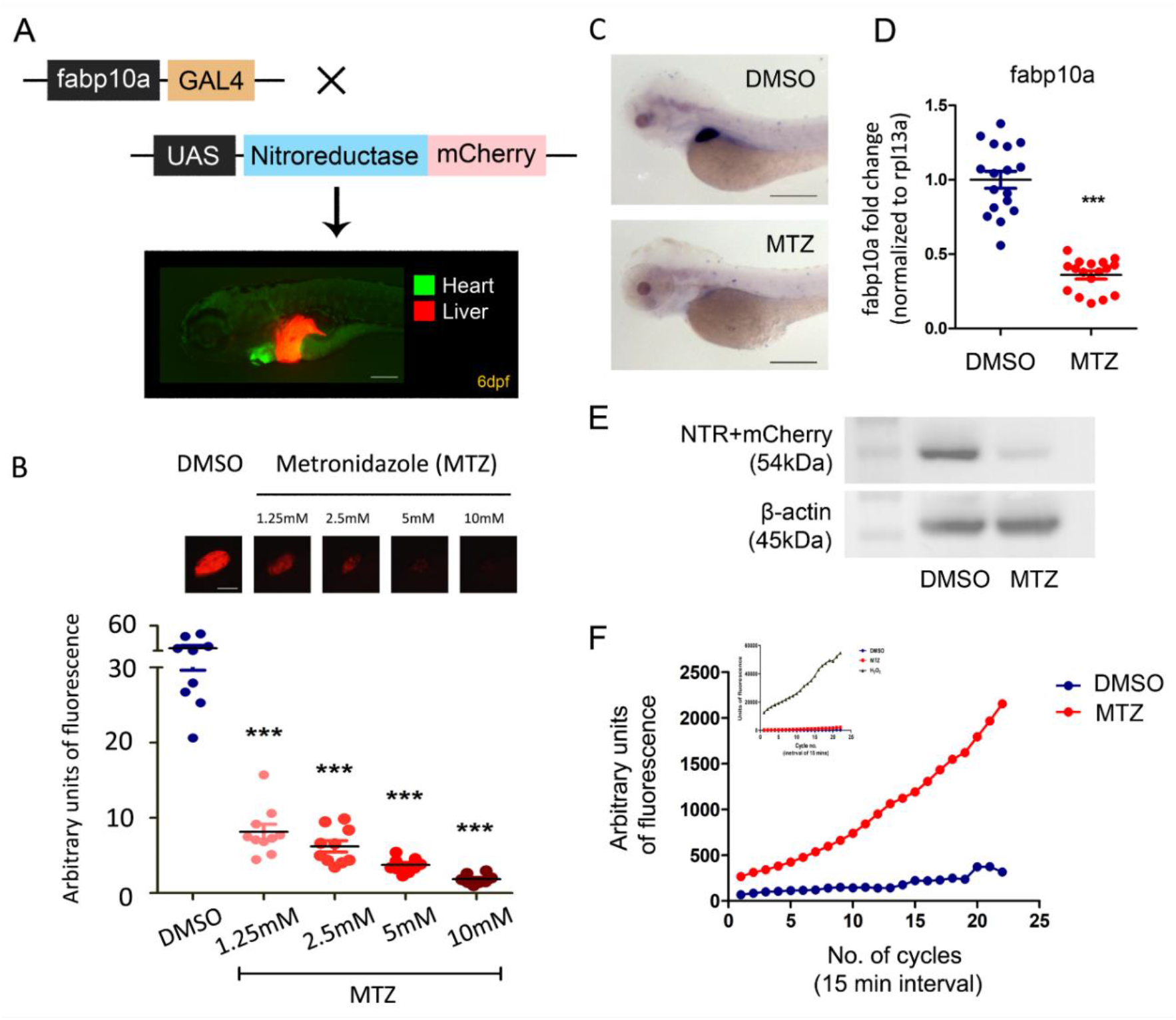
A hepatocyte specific cell ablation system to study liver damage and regeneration in zebrafish. (A) Schematic depiction of double transgenic zebrafish line with a driver line *Tg(fabp10a:GAL4-VP16,myl7:Cerulean)* and effector line: *Tg(UAS:nfsb-mcherry)*. The double transgenic line expresses nitroreductase-mCherry specifically in hepatocytes. Scale bar: 200µm. (B) Upon exposure of pro-drug Metronidazole (MTZ), Nitroreductase converts it into cytotoxic product, leading to hepatocyte ablation (scale bar: 100µ). (C) In situ hybridization (scale bar: 1mm) and (D) quantitative real time measurement of mRNA levels of liver specific fabp10a gene upon MTZ treatment. (E) Western blot for NTR-mCherry fusion protein levels. (F) DCFDA assay for ROS detection in DMSO, MTZ and H_2_O_2_ treated zebrafish embryos. H_2_O_2_ served as a positive control (Black line in the graph inset). *P<0.05, **P<0.01, ***P<0.001, ****P<0.0001.

mCherry fluorescence is being used as a surrogate for liver size in our study. To confirm the damaging effects on the liver we used other markers of hepatocytes. We performed RNA in situ hybridization against fabp10a on 4 dpf larvae treated with 2.5mM MTZ to assess liver size and gene expression. We found an evident expression of fabp10a clearly marking the boundaries of livers of DMSO treated larvae whereas the MTZ treated larvae showed significant reduction in expression (Fig. 1C). We also validated the same using transcript levels of fabp10a, the liver specific marker, by quantitative PCR (Fig. 1D) and observed a significant reduction. This decrease was also associated with decreased mCherry protein expression in whole larval lysates (Fig. 1E).

As reduction of pro-drug Metronidazole has been known to lead to reactive oxygen species [31], we performed DCFDA assay on live larvae treated with DMSO and 2.5mM MTZ. Hydrogen peroxide served as a positive control (Figure 2F, inset). We observed a significant increase in ROS production in MTZ treated larvae as compared to DMSO (Fig. 1F). The experiments validated that mCherry expression can be directly related to liver size and therefore the *nfsb*-NTR system can be utilized for screening of small molecules that could modulate the liver damage and/or regeneration.

**Figure 2:**
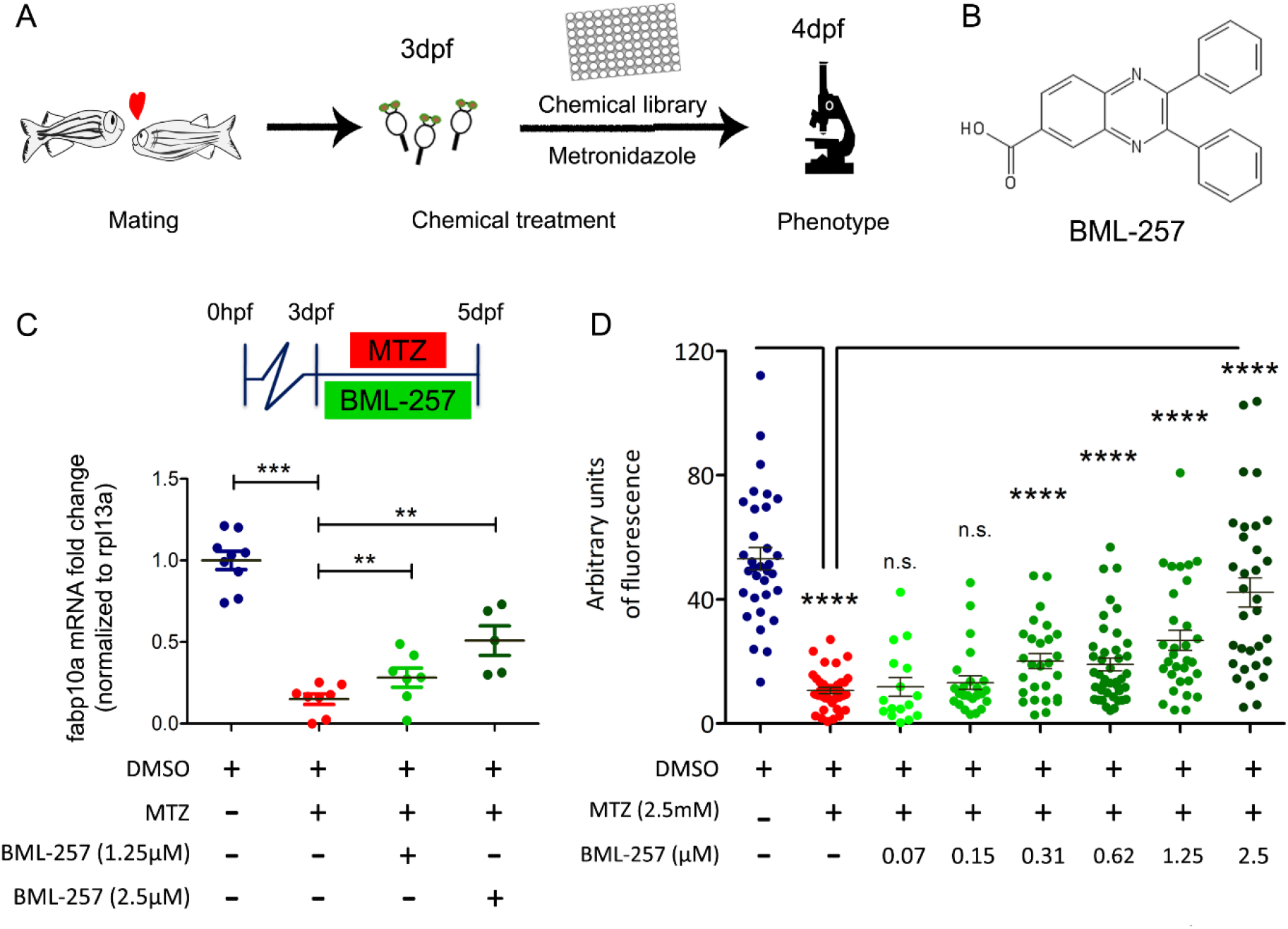
BML-257 shows protective activity against Metronidazole induced liver injury. (A) Screening protocol for small molecule modifiers of MTZ-induced liver injury. (B) Chemical structure of BML-257. (C) BML-257 shows concentration dependent protection from MTZ-induced liver damage in transgenic zebrafish embryos as shown by (C) fabp10a transcript measurements by quantitative real time PCR. Each dot represents a pool of 25 embryos (D) ImageJ based quantification of mCherry fluorescence in liver. *P<0.05, **P<0.01, ***P<0.001, ****P<0.0001.

### BML-257 has a protective effect against MTZ-induced liver damage

To identify compounds that can prevent DILI, we performed a small molecule chemical screen using the *fabp10a-nfsb* transgenic zebrafish larvae. Response to liver damage is a complex process which involves various cell types and signaling pathways. Since protein phosphorylation is one of the critical modifications that modulate numerous pathways, we chose a commercially available kinase inhibitor library (Enzo, SCREEN-WELL®) for our screen that contained 80 molecules. We used mCherry fluorescence as a surrogate to visualize liver damage and regeneration of hepatocytes. We treated 3dpf larvae with 2.5mM MTZ and 10µM of the test drug. After 24 hours of exposure, the larvae were fixed in 4% paraformaldehyde (PFA), the mCherry fluorescence visualized and quantified using ImageJ software (Fig. 2A). Compounds that were toxic at 10µM were titrated down to 2.5µM and we identified BML-257 as a strong candidate against MTZ-induced liver damage (Fig. 2B). The larvae did not show any other morphological deformities at 2.5µM concentration.

To confirm the protective effect of BML-257, we performed quantitative RT-PCR of *fabp10a*. Compared to the MTZ treated livers, co-treatment with BML-257 led to significantly less damage or more liver fluorescence (Fig. 2C). We exposed 3 dpf larvae to 2.5 mM MTZ and co-treated with increasing concentrations of BML-257 (0.078 µM to 2.5 µM). The protective effect of BML-257 appeared to improve with Increasing concentrations (Fig. 2D).

### BML-257 shows prophylactic and regenerative activity in MTZ induced liver injury model of zebrafish

Treatment of *fabp10a-nfsb* with 2.5mM MTZ from 3 dpf to 4 dpf caused specific damage to hepatocytes. We monitored mCherry fluorescence for 4 days post MTZ treatment to visualize and quantify liver damage and regeneration (Fig. 3A). We observed a dramatic reduction in mCherry fluorescence upon 24 hours of MTZ exposure. However, the fluorescence continued to drop for another 24 hours after MTZ was removed from the water indicating the persistence of MTZ in the larvae that continued to cause liver damage. Similar observations have been made in adult *fabp10a-nfsb* liver [19] and in other tissue ablation models using *nfsb*-MTZ system [32]. By imaging and quantification of fluorescence we observed a robust recovery and regeneration of the liver to sizes comparable to undamaged state by 96 hours post damage i.e., 8 dpf (Fig. 3A, 3B). To check if BML-257 had any effect on the damaged liver, we treated MTZ-damaged larvae with BML-257 for 48hours. The red fluorescent liver of the 6 dpf larvae were then imaged and quantified. When compared to the DMSO vehicle control treated animals, BML-257 (2.5 µM) treated livers showed significant increase in size. There was no significant effect of 1.25µM BML-257. This suggests that BML-257 can improve the regenerative abilities of the zebrafish liver.

**Figure 3:**
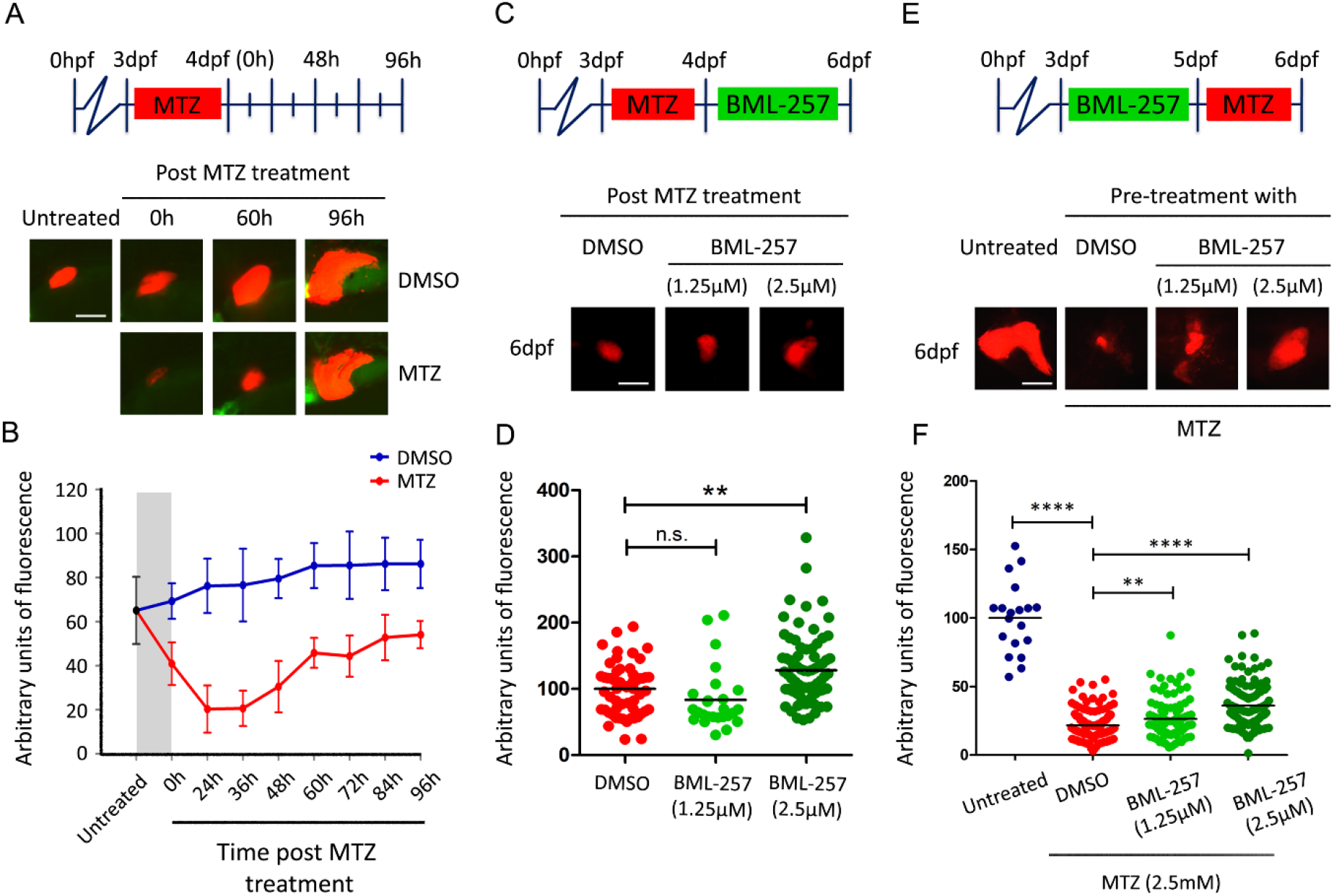
BML-257 shows prophylactic and regenerative activity in MTZ induced liver injury model of zebrafish. (A) mCherry depicts the real time changes in liver size. Upon 24 hrs of MTZ exposure, mCherry fluorescence shows decrease but also shows progressive increase leading to almost full recovery by 96 hours post-MTZ removal. Representative images and (B) ImageJ based quantification of mCherry fluorescence measurement. (C) Assay and representative images for regenerative activity of BML-257 along with (D) ImageJ based quantification of mCherry fluorescence. (E) Assay for prophylactic activity of BML-257. Zebrafish embryos first treated with BML-257 for 48 hours followed by MTZ exposure for 24 hours. Representative images and (F) ImageJ based quantification of mCherry fluorescence. A minimum of 30 embryos have been used for each experiment and all experiments are repeated at least 3 times. Scale bar: 100µm, *P<0.05, **P<0.01, ***P<0.001, ****P<0.0001.

In order to check if the regeneration that we observed is not a result of mere proliferation of hepatocytes, we tested if BML-257 has proliferative activity. We performed the treatment of DMSO and BML-257 in healthy 3dpf larvae. After treatment of 24 hours, the larvae were imaged to detect the increase in mCherry fluorescence, if any. No observable differences were identified (Data not shown) indicating that BML-257 doesn’t have proliferative function by itself.

Since BML-257 appeared to improve recovery from damage as well as reduce the damage, it suggested that the compound may have beneficial effects on the liver prophylactically. We therefore treated 3 dpf larvae with BML-257 for two days and then induced liver damage for 24 hours with MTZ. To our surprise, we found that compared to DMSO pre-treated larvae, BML-257 pre-treated larvae had significantly less damage to the liver. 2.5 µM BML-257 had stronger effects than 1.25 µM.

### BML-257 shows hepatoprotective activity in various clinically relevant toxicological models of liver injury

Acetaminophen overdose is a major cause of DILI in many countries [33, 34]. Isoniazid, another hepatotoxic drug, is also one of the frontline drugs for Tuberculosis treatment [35-37]. We created liver damage models with these hepatotoxic drugs in zebrafish larvae. 24 hours of exposure to 15mM acetaminophen (APAP) caused a marked reduction in liver fluorescence in 4 dpf *fabp10a-nfsb* zebrafish larvae. Similarly, strong effect was seen in larvae treated with 10mM isoniazid (INH). We co-treated larvae with (a) APAP and BML-257 and with (b) INH and BML-257 then quantified the mCherry fluorescence in 4 dpf larvae. Animals co-treated with 2.5mM BML-257 and 15mM APAP showed a significant improvement in liver fluorescence compared to APAP and DMSO treated fish. 1.25µM BML-257 did not have a significant effect on the liver damage caused by APAP. On the other hand, both 1.25µM and 2.5µM BML-257 showed significant increase in liver fluorescence when co-treated with 10mM INH compared to fishes treated with INH and DMSO (Fig. 4B, C).

**Figure 4:**
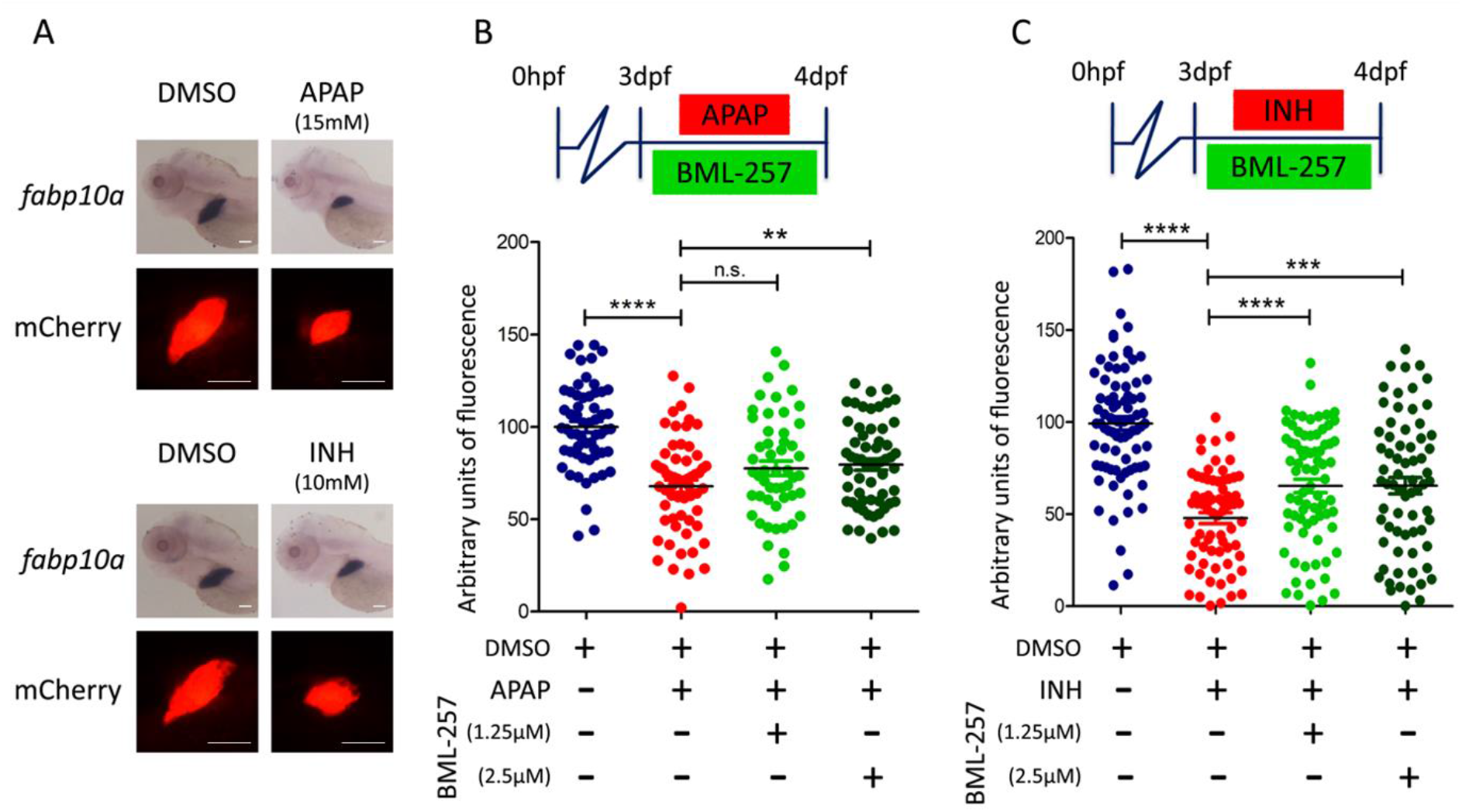
BML-257 shows hepatoprotective activity in various clinically relevant toxicological models of liver injury. (A) Drug induced liver injury (DILI) models of Acetaminophen and Isoniazid as shown by in-situ hybridization of fabp10a riboprobe as well as mCherry fluorescence in liver. Hepatoprotective activity of BML-257 was assessed in both (B) Acetaminophen and (C) Isoniazid induced liver injury models. Zebrafish embryos were co-exposed to BML-257 and offending drug for 24 hours. Representative images and ImageJ-based quantification of mCherry fluorescence is plotted. A minimum of 25 embryos have been used for each experiment and all experiments are repeated at least 3 times. Scale bar: 100µm, **P<0.01, ***P<0.001, ****P<0.0001.

## Conclusion

Molecules that can protect the liver from damage or augment the liver’s natural regeneration potential are valuable. Various plant natural extracts have long been used for their hepatoprotective activities. Various general liver wellness preparations are available in traditional medicines such as Ayurveda. However, in most such cases the active components have not been identified thus making elucidation of the mechanism of action or understanding the deleterious side effects difficult. In this study, we have used a MTZ based hepatocyte ablation system in zebrafish to screen for molecules that can protect the liver from hepatotoxic insults. From a library of kinase inhibitors, we identified a MAPK inhibitor, BML-257 that shows robust hepatoprotective activity, both in a prophylactic protection assay as well as when administered with the damaging agent, MTZ. This protective activity of BML-257 is strong even when we change the type and agent of liver damage. Acetaminophen (APAP) causes hepatocellular death by GSH depletion, superoxide and peroxynitrite formation, protein adduct formation and lysosomal iron uptake into mitochondria. As one of the most widely used analgesics, APAP is prone to overuse causing liver damage leading to hospitalization and death. Isoniazid (INH) is one of four drugs given as part of the DOT therapy for eradicating TB.

For the globally devastating pandemic of Tuberculosis treatment INH is indispensable. However, INH forms covalent adducts to liver macromolecules thereby leading to reactive metabolites and immune response. The associated hepatotoxicity of INH weakens an already compromised system of the TB patient. BML-257 showed strong protection from liver damage when administered with either APAP or INH in the zebrafish larval model. We also found that BML-257 was moderately effective in improving the regeneration of liver after the hepatocyte cell death caused by MTZ treatment.

## Acknowledgements

We thank Shashi Ranjan and Narendra Kumar for the maintenance of the zebrafish facility. We thank A. Agrawal, R. Gokhale, M. Datta for discussion and critical inputs. Fabp10a plasmid was a kind gift from Yann Gilbert and Paula Fraenkel. Anusha Krishnan helped with the mCherry antibody.

## Author contributions

CS conceived, supervised and secured funding for the project. CS and UJ designed experiments as well as wrote the manuscript. UJ characterized the *fabp10a-nfsb* transgenic line, performed drug screening and all other assays. SB assisted in drug screening, BML-257 dosage standardization and data analysis. LL and NB standardized APAP and INH dosage.

